# Angiotensin II and cAMP signaling pathways regulate mitochondrial biogenesis and activity in human adrenocortical cells

**DOI:** 10.64898/2026.05.06.723032

**Authors:** Matías A. Belluno, Fabrizio G. Arona, Katia E. Helfenberger, Martina A. Rodrigo, María Mercedes Mori Sequeiros Garcia, Paula M. Maloberti, Yanina Benzo, Cecilia Poderoso

## Abstract

Mitochondrial homeostasis, governed by the balance between biogenesis and mitophagy, is essential for steroidogenesis in adrenocortical cells. While the requirement of active mitochondria for steroid synthesis is well-established, the hormonal regulation of genes governing mitochondrial function remains poorly understood. This study investigated whether angiotensin II (Ang II) and the cAMP/PKA pathway modulate the expression of key regulatory factors involved in mitochondrial biogenesis and redox status in the human adrenocortical H295R cell line.

Using real-time qPCR and Western blot, we show that Ang II and 8Br-cAMP —a permeant analogue of cAMP— modulate NRF-1, Nrf2, UCP2, and ANT1 impacting on mitochondrial biogenesis, antioxidant defense, and respiratory activity. These molecular changes correlated with increased mitochondrial membrane polarization, as confirmed by MitoTracker red staining. Interestingly, Ang II stimulation promoted a time-dependent increase in TFAM levels, a key transcription factor in mitochondria, which correlates with the increase in mitochondrial DNA (mtDNA) content. The rate of oxygen consumption (OCR) and mitochondrial parameters were determined, with results showing that Ang II led to a significant increase in basal and maximum respiration, ATP production, and proton leak. These findings suggest that hormone stimulation favors mitochondrial activity, thereby enhancing the bioenergetic capacity of adrenocortical cells. Furthermore, treatment with the uncoupler CCCP triggered a retrograde signaling response, upregulating nuclear-encoded mitochondrial genes to counteract mitochondrial membrane depolarization.

Our findings demonstrate for the first time that hormonal signals directly modulate the mitochondrial genetic program in H295R human adrenocortical cells, optimizing the bioenergetic platform required for efficient steroidogenic function.

## Introduction

Mitochondrial biogenesis is a complex, multi-step process defined as the growth and division of pre-existing mitochondria, a mechanism critical to cellular homeostasis and metabolic adaptation [1,2]. This process requires fine-tuned coordination between the mitochondrial and nuclear genomes, as over 98% of mitochondrial proteins are encoded in the nucleus. Biogenesis is primarily orchestrated by the peroxisome proliferator-activated receptor gamma coactivator 1 (PGC-1) family (PGC-1α, PGC-1β, and PRC), which act as master transcriptional co-activators [3]. In particular, PGC-1α is often described as a master regulator of mitochondrial biogenesis as well as a central player in regulating the antioxidant defense [4] and various physiological signals, such as thermogenesis, glucose metabolism, and steroidogenesis [5–7]. PGC-1α is crucial to initiate the recruitment of a well-known family of nuclear respiratory factors (NRF), including NRF-1 and NRF-2 (or GABPA), and acts as a co-activator of estrogen-related receptor α (ERRα) [8–10]. These transcription factors subsequently activate the expression of the mitochondrial transcription factor A (TFAM) and mitochondrial transcription factor B2 (TFBM), which act as final effectors by binding to the mitochondrial DNA (mtDNA) promoter to drive transcription and replication [11]. Mitochondrial biogenesis increases the mitochondrial mass and promotes inner membrane remodeling, leading to greater cristae density where electron transport chain (ETC) complexes are localized. Then, oxidative phosphorylation (OXPHOS) and respiratory capacity increases [12].

The regulation of this mitochondrial system is highly responsive to hormonal signaling. Angiotensin II (Ang II) stimulates aldosterone biosynthesis in the adrenal zona glomerulosa, while the ACTH/cAMP/PKA axis regulates glucocorticoid production in the zona fasciculata, with steroidogenesis being intrinsically linked to mitochondrial activity [13,14]. Because Leydig cell steroidogenesis depends on energized, polarized, and actively respiring mitochondria, alterations in mitochondrial function may influence steroid biosynthesis [15].

Ang II signaling leads to an increase in reactive oxygen species (ROS) production [16] and this increase in ROS acts as an important signal modulating the nuclear factor-erythroid-derived 2-like (Nrf2, also known as 2NFE2L2), a master regulator of the cellular antioxidant response [17]. Nrf2 has been shown to increase the expression of the NRF-1 and PGC-1α genes —thus promoting mitochondrial biogenesis [17,18]— and regulates the balance between oxidative phosphorylation and the cellular redox state.

Mitochondrial homeostasis is further preserved by physiological proton leaks mediated by mitochondrial uncoupling proteins (UCPs) and adenine nucleotide translocases (ANTs) [19]. These proteins are not merely involved in energy balance but act as sensors of mitochondrial stress. Particularly, UCP type 2 (UCP2) mediates mild proton leak across the inner mitochondrial membrane, helping to regulate mitochondrial membrane potential (ΔΨ*m*) and limit excessive ROS production [20]. ANT isoform 1 (ANT1) regulates basal proton conductance and mediates ADP/ATP exchange, thus playing a crucial role in mitochondrial integrity. When mitochondrial function is compromised —either for endogenous or exogenous reasons— cells initiate retrograde signaling to communicate the organelle’s status to the nucleus [21]. This adaptive mechanism is initiated by mitochondrial stress, such as the dissipation of the electrochemical gradient by agents like the uncoupler carbonyl cyanide m-chlorophenyl hydrazone (CCCP), a protonophore that strongly promotes a decrease in ΔΨ*m* and thus blocks ATP synthase activity. Mitochondrial dysfunction triggers signal release including elevated Ca^2+^ fluxes, ROS, and altered AMP/ATP ratios, which in turn induces the expression of NRF-1/2 and PGC-1α and the promotion of cellular proliferation, anti-apoptotic signals, and mitophagy, among others [22].

The aim of this study was to determine whether hormone stimulation regulates the expression of nuclear/mitochondrial transcription factors, antioxidant signaling, and mitochondrial respiratory activity. We assessed if mitochondrial dysfunction triggers a compensatory retrograde signaling response in the human adrenocortical cell line H295R.

Our findings demonstrate that Ang II and cAMP regulate mitochondrial biogenesis and bioenergetic function in H295R cells. Moreover, CCCP-induced depolarization activates retrograde signaling, supporting functional mitochondria-to-nucleus communication. Together, these results provide the first evidence that hormonal signaling coordinates mitochondrial adaptation to sustain steroidogenic function.

## Materials & Methods

### H295R cell line and treatments

The NCI-H295R cell line is a clonal strain of human adrenal carcinoma. The cell line was from the American Type Culture Collection (ATCC, Manassas, VA, US) and was handled as originally described [23]. Trypsin-EDTA and antibiotics were acquired from Gibco-Life Technologies (Gaithersburg, MD, US). Plasticware for cell culture was purchased from Corning-Costar (Corning, NY, US). H295R cells were maintained in DMEM/Ham’s F12 1:1 medium containing 1.1 g/liter NaHCO_3_, 20 mM HEPES, 200 IU/ml penicillin, 200 μg/ml streptomycin sulfate, supplemented with 5% Cosmic Calf® serum (HyClone, Cytiva, Chicago, IL, US) and 1% penicillin-streptomycin at 37 °C in a 5% CO_2_ humidified atmosphere. Flasks and multi-well plates were maintained at 37 °C in a humidified atmosphere containing 5% CO_2_. For experiments, cells (until passage number 15) were serum-starved for 48 hours and after that, they were incubated with 100 nM Ang II or 1 mM 8Br-cAMP (Sigma-Aldrich, Burlington, MA, US) in serum-free medium, for different times. The choice of 100 nM Ang II was based on extensive literature [24–27]. When applicable, cells were pre-incubated 30 min with 5 μM CCCP (Sigma-Aldrich) to assess ΔΨ*m*, as previously reported [26,28,29]. At the end of treatments, monolayers were washed with phosphate-buffered saline (PBS) 1x and harvested for different purposes.

### Protein lysates and nuclear fractionation

Total protein lysates were obtained using a buffer containing 20 mM Tris-HCl, 1 mM EDTA, 1 mM EGTA, 125 mM NaCl, 1% Triton X-100, and a cocktail of protease and phosphatase inhibitors (leupeptin, pepstatin A, PMSF, NaF, and β-glycerophosphate). Cells were harvested mechanically and centrifuged (1,000 x g, 10 min, 4 °C). For nuclear fractions, cells were collected in cold PBS, resuspended in a low-salt buffer (10 mM HEPES, 10 mM KCl, 0.1 mM EDTA/EGTA, DTT, and protease inhibitors), and incubated on ice for 15 min. Following the addition of 10% NP-40 and a high-salt buffer (20 mM HEPES, 25% glycerol, 0.4 M NaCl), samples were homogenized using a tuberculin syringe and centrifuged (13,000 x g, 15 min, 4 °C). The supernatant was collected as the soluble nuclear fraction [30].

### Isolation of mitochondria

Mitochondria were isolated as previously described [31]. Briefly, cell cultures were washed with PBS, scraped in 10 mM Tris-HCl (pH 7.4), 250 mM sucrose, 0.1 mM EDTA, 10 μM leupeptin, 1 μM pepstatin A, and 1 mM EGTA (buffer A), homogenized with a pellet pestle motor homogenizer (Kimble Kontes), and centrifuged at 1,000*x*g for 10 min. The supernatant was centrifuged at 18,000*x*g for 20 min and rendered a mitochondrial pellet that was resuspended in buffer A. Control fraction markers were performed as in our previous work [25].

### Western blot

Mitochondrial and nuclear proteins were separated on 12% or 10% SDS-PAGE and electrotransferred to PVDF membranes (Bio-Rad Laboratories, Hercules, CA, US) as previously described [27] The proteins transferred were visualized using a Ponceau S staining solution containing 0.2% Ponceau S in 1% acetic acid. Membranes were then incubated with 1% bovine serum albumin (BSA) in 500 mM NaCl, 20 mM Tris-HCl pH 7.5, and 0.5 % Tween 20 (T-TBS) for 2 h at room temperature, with gentle shaking. The membranes were then rinsed twice with T-TBS and incubated overnight at 4 °C with 1:2000 dilutions of primary antibodies mouse monoclonal anti-NRF-1 (RRID:AB_1126766), anti-PCNA (RRID:AB_628109), anti-TFAM (RRID:AB_10610743) (Santa Cruz Biotechnology Inc, Dallas, TX, US) and rabbit monoclonal anti-succinate dehydrogenase subunit A (SDHA) (RRID:AB_2750900) (Cell Signaling Technologies, Danvers, Massachusetts, US). Bound antibodies were visualized by incubation with 1:5000 horseradish peroxidase-conjugated secondary goat anti-mouse (RRID:AB_11125547) or 1:5000 anti-rabbit antibodies (RRID:AB_11125142) (Bio-Rad Laboratories Inc.) and detected by chemiluminescence (BioLumina, Kalium Tech, Buenos Aires, Argentina). The immunoblots were then quantified using Gel Pro Analyzer software.

### RNA extraction and real-time PCR

For real-time quantitative PCR (qPCR), total RNA was isolated using Tri Reagent following the manufacturer’s instructions (Molecular Research Center Inc., Cincinnati, OH, US). Extracted RNA was deoxyribonuclease-treated using RNAse-free DNase RQ1 (Promega, Madison, WI, USA). Reverse transcription was performed using total RNA (2 μg) and M-MLV Reverse Transcriptase (Promega). The expression of NRF-1 (forward primer: 5’-GTACAAGAGCATGATCCTGGA-3’, reverse primer: 5’-GCTCTTCTGTGCGGACATC-3’), Nrf2 (forward primer: 5’-CAGCGACGGAAAGAGTATGA-3’, reverse primer: 5’-TGGGCAACCTGGGAGTAG-3’), UCP2 (forward primer: 5’-TGGTCGGAGATACCAAAGCAC-3’, reverse primer: 5’-GCTCAGCACAGTTGACAATGGC-3’), and ANT1 (forward primer: 5’-TCAACGTCTCTGTCCAAGGC-3’, reverse primer: 5’-GTCAACTGTCCCCGTGTACA-3’) were assessed by real time qPCR (Mic PCR instrument, Molecular Biosystems, San Diego, CA, US) using the FastStart Universal SYBR Green Master (ROX) (Roche, Mannheim, Germany). The reaction conditions were: one cycle at 95 °C for 5 min, followed by 40 cycles at 95 °C for 15 s, 60 °C for 30 s, and 72 °C for 30 s. Gene mRNA expression levels were normalized to human cyclophilin (forward primer: 5’-TGGCAAGTCCATCTATGGGGA-3’, reverse primer: 5’-ACTTATTCGAGTTGTCCAACAGTCAGCA-3’). Real-time PCR data were analyzed by calculating the 2^-ΔΔCt^ value (comparative Ct method) for each experimental sample (relative gene expression).

### Mitochondrial activity assay

H295R cells were seeded on poly-L-lysine-coated glass coverslips. Following stimulation, cells were incubated with 150 nM MitoTracker Deep Red FM (Invitrogen, Waltham, MA, US) for 45 min at 37°C in the dark. This fluorescent dye accumulates in mitochondria with active respiration and high membrane potential (excitation 581 nm/emission 644 nm). Cells were fixed in 4% paraformaldehyde/5% sucrose, counterstained with 4′,6-diamidino-2-phenylindole (DAPI), mounted with DAKO mounting medium (Agilent Technologies, Santa Clara, CA, US) and analyzed using a Nikon ECLIPSE E200 MV epifluorescence microscope (40X magnification). Images were processed using FIJI/ImageJ software, measuring fluorescence intensity across defined regions of interest (ROI; 15–20 cells per ROI, n=3). Images were not manipulated for analysis.

### Bioenergetics analysis

The oxygen consumption rate (OCR) of H295R cells was measured using the Seahorse XFe96 Extracellular Flux Analyzer (Agilent Technologies, Santa Clara, CA, US). Briefly, 100,000 cells per well were seeded in 96-well Seahorse plates (Seahorse Bioscience, North Billerica, MA, US) pre-coated with polyethylenimine, using DMEM/F12 medium supplemented with 5% Cosmic serum and 1X ITS+1. On the day of the assay, cells were treated with Ang II for the times indicated. Subsequently, the culture medium was replaced with assay medium (3.5 mM KCl, 120 mM NaCl, 1.3 mM CaCl_2_, 0.4 mM KH_2_PO_4_, 1.2 mM Na_2_SO_4_, 2 mM MgSO_4_, 15 mM D-glucose with or without 10 mM pyruvate, and 4 mg/ml BSA, pH 7.4). Plates were incubated at 37°C for 20 min before being loaded into the Seahorse XFe96 analyzer. All assays were conducted at 37 °C. To assess mitochondrial respiratory parameters, the Seahorse XF Cell Mito Stress Test was employed, using the following inhibitors and uncouplers: rotenone/antimycin A, carbonyl cyanide-4-(trifluoromethoxy)phenylhydrazone (FCCP), and oligomycin. Drugs were diluted to final working concentrations of 0.5 μM for rotenone/antimycin A, 0.75 μM for FCCP, and 1.5 μM for oligomycin, and loaded into the sensor cartridge ports according to the manufacturer’s injection protocol. Following the assay, total cellular protein concentration was determined using the Bradford assay. OCR was measured as picomole O_2_ per minute (pmol O_2_/min). Respiratory parameters like basal, maximal, and ATP-linked respiration, proton leak, efficiency coupling, and spare respiratory capacity were calculated based on OCR data obtained in the Mito Stress Tests using Seahorse Wave software for XF analyzers (Agilent). OCR data were normalized to protein content. The experiments were performed at least three times independently.

### mtDNA analysis

mtDNA content was assessed by measuring mtDNA copy number markers through qPCR. Total DNA was extracted from H295R cells previously subjected to various Ang II stimulation times. Using total DNA as a template, qPCR was performed to simultaneously quantify specific nuclear and mitochondrial DNA markers. The expression of β-actin (forward primer: 5’-CTGTGGCATCCACGAAACTA -3’, reverse primer: 5’-AGTACTTGCGCTCAGGAGGA-3’), mitochondrial *b* cytochrome (mtCytB) (forward primer: 5’-AGACAGTCCCACCCTCACAC -3’, reverse primer: 5’-GGTGATTCCTAGGGGGTTGT -3’), mitochondrial non-nuclear segments (mtNonNUMT) (forward primer: 5’-CCCCTCCCACTCCCATACTA -3’, reverse primer: 5’-CCAGCTATCACCAAGCTCGT -3’), and mitochondrial DLoop (mtDLoop3) (forward primer: 5’-CTAGAAACCCCGAAACCAAA -3’, reverse primer: 5’-GGGCGGGGGTTGTATTGAT -3’) was assessed, and mitochondrial signals were normalized against nuclear markers to determine the relative mtDNA content for each experimental condition.

### Protein quantification and statistical analysis

Protein concentration was determined through the Bradford assay [32] using BSA as a standard. Results are expressed as mean ± standard deviation (SD). Statistical significance was determined by one-way analysis of variance (ANOVA) followed by Tukey’s post-hoc test for multiple comparisons. Differences were considered statistically significant at *p < 0.05, **p < 0.01, and ***p < 0.001. All statistical analyses were performed using GraphPad Prism software (version 9.5.1, GraphPad Software, San Diego, CA, US).

## Results

### The expression of genes that regulate mitochondrial biogenesis, redox status, and mitochondrial activity is modulated by Ang II

We investigated whether Ang II stimulation modulates NRF-1 expression, given that mitochondria are essential for steroidogenesis in adrenocortical cells. H295R cells were stimulated with Ang II for the times indicated, and real-time qPCR was performed to analyze NRF-1 expression. Our results demonstrate that Ang II promotes a time-dependent increase in NRF-1 mRNA expression, with a peak at 2 h post-stimulation and a subsequent decline (Figure 1, panel A). In addition to mRNA expression, protein expression analyses revealed an increase in NRF-1 within the nuclear fraction upon Ang II treatment (Figure 1, panel B). The temporal activation profile observed for protein levels was consistent with the kinetics of NRF-1 mRNA upregulation. Furthermore, the protein levels of TFAM —which is positively regulated upstream by NRF-1— were increased by Ang II in a time-dependent manner (Figure 1, panel C). These findings suggest that the hormonally regulated increase in NRF-1 may exert a functional effect on TFAM transcription, thereby modulating mitochondrial activity and biogenesis in H295R adrenocortical cells.

**Figure 1.**
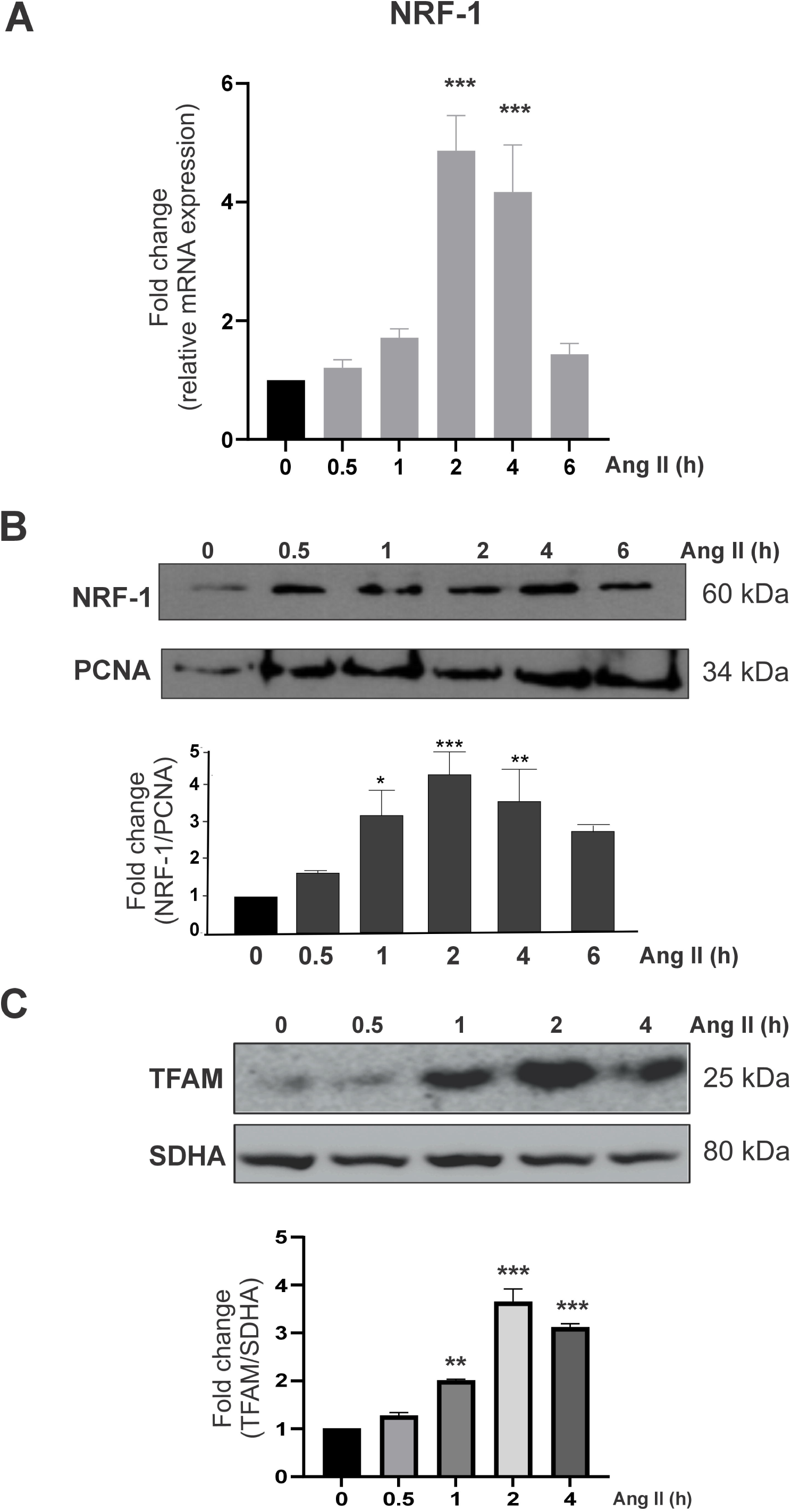
Ang II modulates mitochondrial biogenesis factors expression. H295R cells were serum-starved for 24 h and subsequently stimulated with angiotensin II (Ang II, 100 nM) for the times indicated (hours, h). Total RNA was isolated and subjected to real-time qPCR using specific primers. Cyclophilin was used as an internal control. qPCR data were analyzed using the 2^-ΔΔCt^ method (comparative Ct method) for each experimental sample. (A) Relative NRF-1 expression levels (fold change): *** p < 0.001 vs. control cells (Ang II 0 h). (B) Nuclear and (C) mitochondrial proteins were analyzed by Western blot. Membranes were sequentially incubated with (B) anti-NRF-1 and anti-PCNA and (C) anti-TFAM and anti-SDHA antibodies. Representative images are shown from 3 independent experiments (n=3). Optical density for each band was taken, and control (without Ang II) sample intensity was arbitrarily defined as 1. Protein levels were relativized to respective controls (fold change): ***p<0.001; **p<0.01; *p<0.05 vs respective control (Ang II 0 h). Results are expressed as mean ± SD of three independent experiments. ANOVA followed by Tukey’s post hoc test.

We subsequently evaluated Nrf2 expression following Ang II treatment. As shown in Figure 2, Ang II signaling induced a slight decrease in Nrf2 mRNA expression within the first 30 min, followed by a significant increase sustained from 2 to 6 h of stimulation (Figure 2).

**Figure 2.**
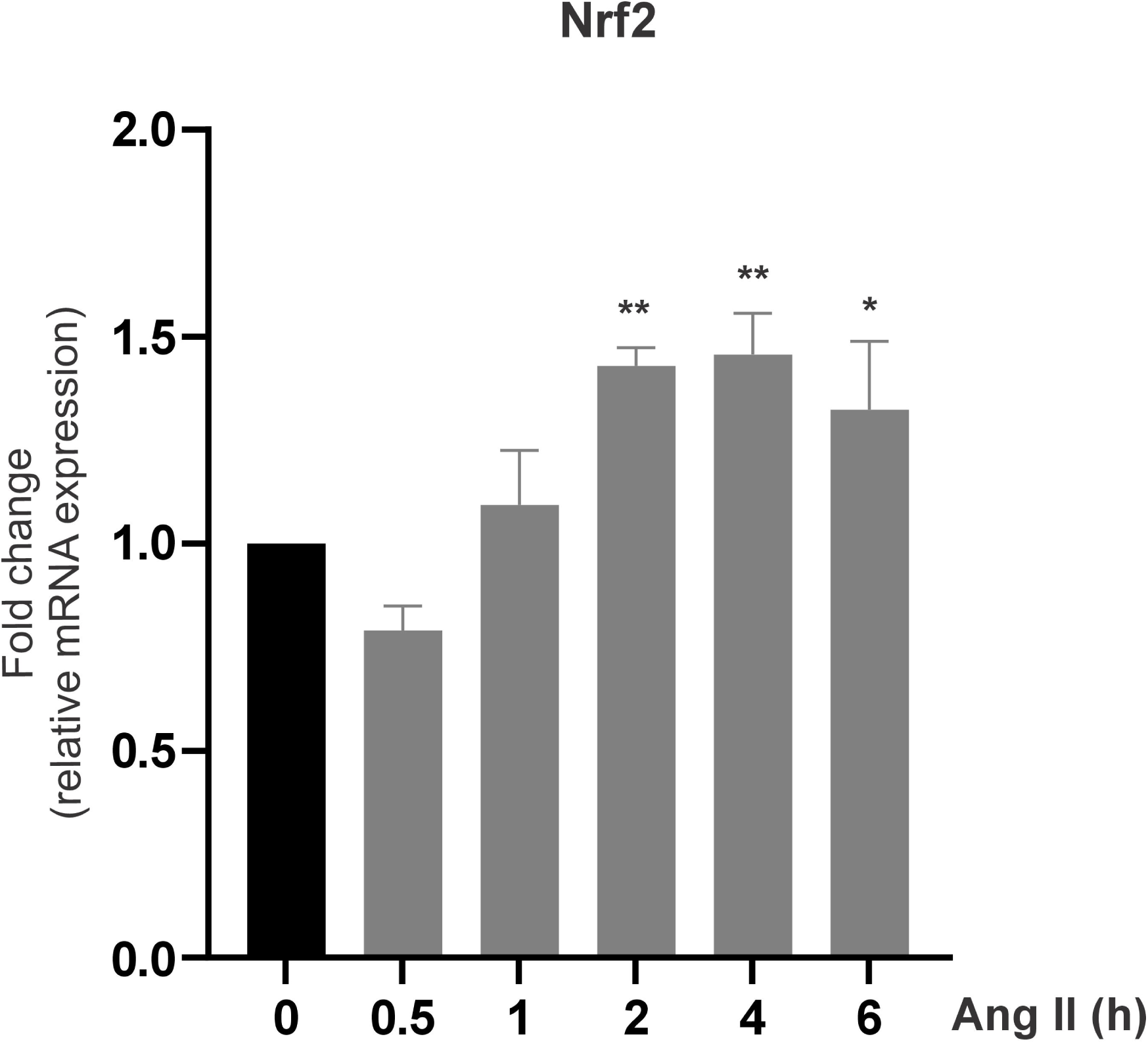
Ang II modulates Nrf2 mRNA levels in a time-dependent manner. H295R cells were serum-starved for 24 h and subsequently stimulated with angiotensin II (Ang II, 100 nM) for the times indicated (hours, h). Total RNA was isolated, and real-time qPCR was performed using specific primers. Cyclophilin was used as an internal control. qPCR data were analyzed using the 2^-ΔΔCt^ method (comparative Ct method) for each experimental sample. Representative images are shown from 3 independent experiments (n=3). Relative Nrf2 expression levels (fold change): *p<0.05; **p<0.01 vs. control cells (Ang II 0 h). Results are expressed as mean ± SD of three independent experiments. ANOVA followed by Tukey’s post hoc test.

Given that UCP2 is ubiquitously expressed and regulates ROS levels, and that Ang II modulates ROS production in H295R adrenocortical cells, we aimed to investigate UCP2 expression under Ang II stimulation. Ang II regulated UCP2 expression with a mild decrease up to 2 h but significant peak of mRNA expression at 4 h of Ang II exposure (Figure 3, panel A).

**Figure 3.**
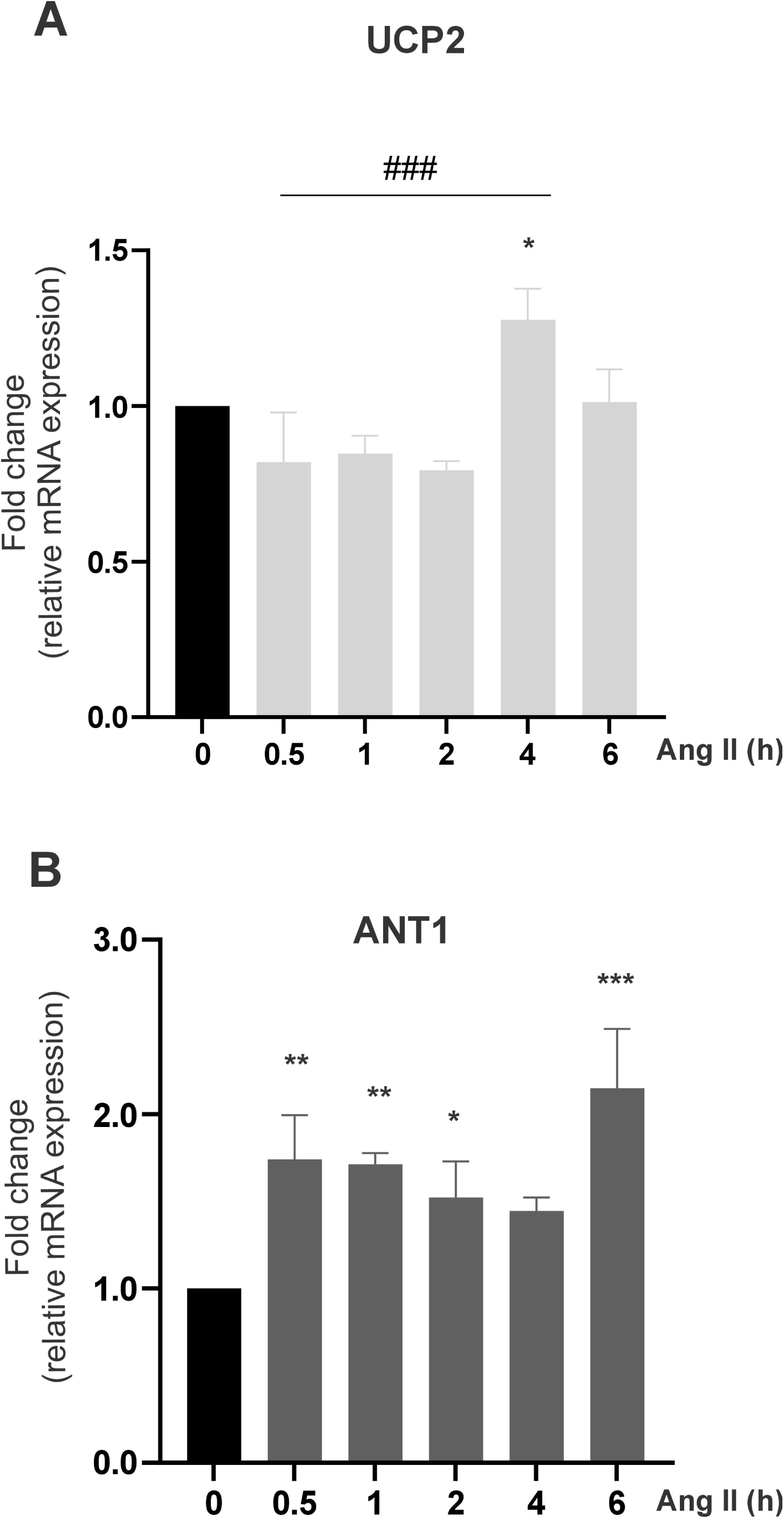
UCP2 and ANT1 expression is regulated by Ang II. H295R cells were serum-starved for 24 h and subsequently stimulated with angiotensin II (Ang II, 100 nM) for the times indicated (hours, h). Total RNA was isolated and subjected to real-time qPCR using specific primers. Cyclophilin was used as an internal control. qPCR data were analyzed using the 2^-ΔΔCt^ method (comparative Ct method) for each experimental sample. Relative (A) UCP2 and (B) ANT1 expression levels (fold change): *p<0.05; **p<0.01; *** p<0.001 vs. control cells (Ang II 0 h); ^###^p<0.001 vs. 0.5, 1 and 2 h Ang II. Data were analyzed by ANOVA followed by Tukey’s post-hoc test.

The physiological uncoupling effect of UCP2 might be initially inhibited to allow Ang II-induced ROS to promote stimulatory effects on steroid production.

As ANT1 is known to regulate processes such as proton leak, electron leak, and proton slip [30], we conducted real-time qPCR to assess ANT1 mRNA levels following hormonal stimulation. Results showed a significant increase as early as 30 minutes post-stimulation with Ang II, with a peak at 6 h, showing a biphasic effect of hormonal signaling on ANT1 expression (Figure 3, panel B).

### Ang II positively regulates mitochondrial activity

Mitochondrial activity is represented by the oxygen consumption rate, ATP synthesis, and the polarization of the inner mitochondrial membrane, which is driven by the proton-motive force and ΔΨ*m*.

Fluorescence microscopy studies determined that highly active mitochondria exhibit a stronger red fluorescent signal; in agreement, we observed that Ang II stimulation increased fluorescent signal intensity over time, which indicates that hormonal signaling promotes greater mitochondrial bioenergetic activity (Figure 4, panels A–D). Quantification assays confirmed a progressive trend throughout the stimulation period. The right-hand column (CCCP treatment) served as a negative control, showing a marked decrease in the fluorescent signal. Fluorescence intensity was quantified relative to the number of cells (nuclei) in each region of interest (ROI) analyzed (Figure 4, panel E).

**Figure 4.**
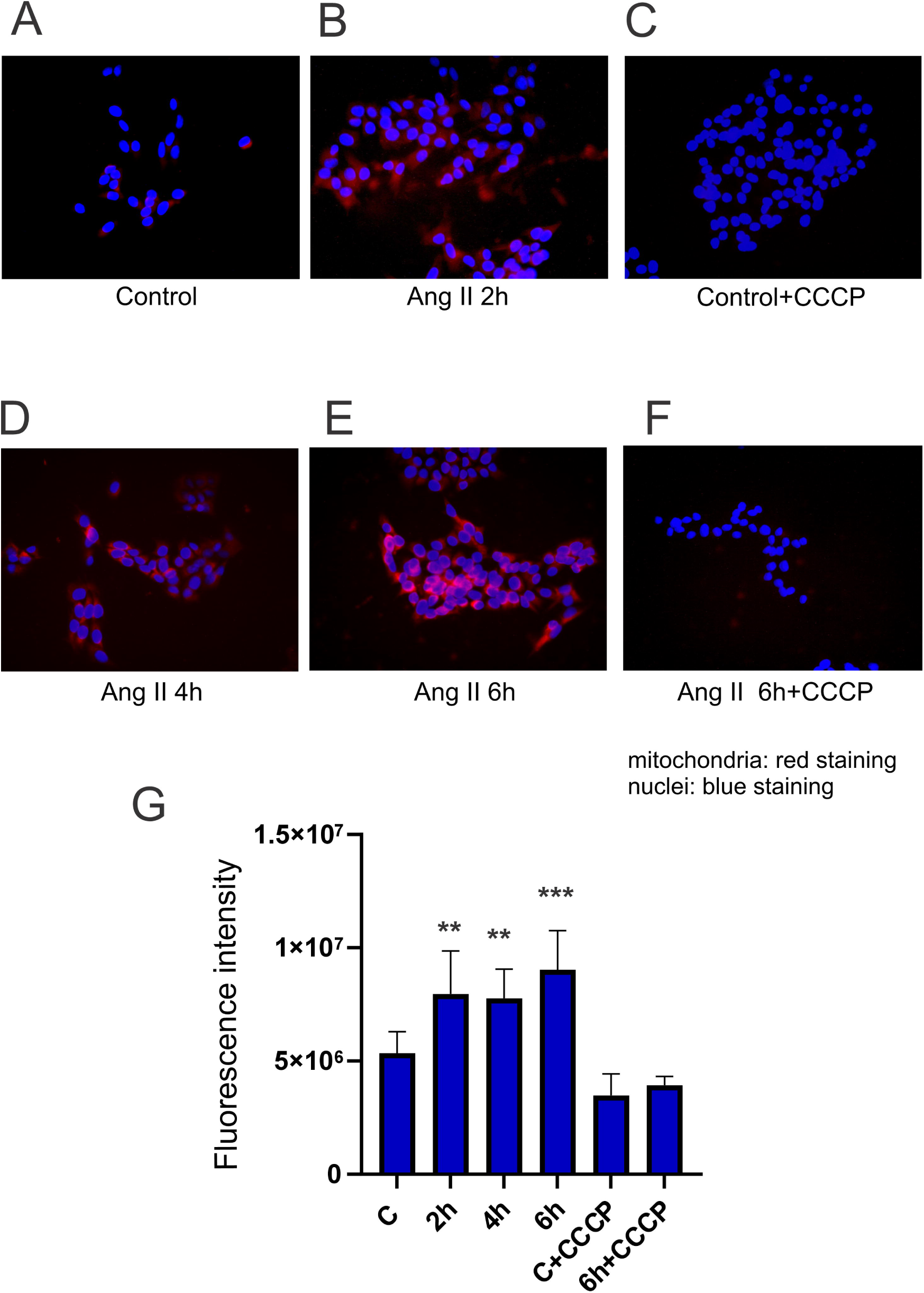
Ang II stimulation promotes mitochondrial activity. H295R cells were grown on poly-L-lysine glass coverslips, serum-starved for 24 h, and subsequently incubated with CCCP (5 μM) and/or DMSO (0.1%) as vehicle in the corresponding wells (F), followed by stimulation with or without angiotensin II (Ang II, 100 nM) for the times indicated (hours, h) (A-F). Afterwards, cells were incubated with 150 nM MitoTracker Red (MTR) at 37°C in fresh medium for 45 min and samples were processed to be analyzed using a Nikon fluorescence microscope (40x magnification). (G) In the bar graph, differences in red signal intensity (mitochondrial signal) are expressed as fluorescence intensity (FIJI software) relative to the number of cells within the same ROI, (containing 15–20 cells/ROI in all cases). Red (MTR) and ultraviolet (DAPI) channels were studied. Representative images are shown of 10 images analyzed from 3 independent experiments (n=3). Images were obtained in all cases with the same exposure, the background was discarded, and the intensity of the red fluorescent signal was analyzed with FIJI software. **p<0.01; ***p<001 vs control (Ang II 0 h). Scale bar: 20 µm for every image. Data were analyzed by ANOVA followed by Tukey’s post-hoc test.

### Mitochondrial DNA content is modulated by Ang II

In view of our previous findings showing that Ang II stimulation promotes mitochondrial fusion in a time-dependent manner and positively modulates MFN2 [27], we next aimed to evaluate whether Ang II incubation also affects the mitochondrial mass and other mitochondrial parameters. To this end, the relative quantity of mtDNA was analyzed with respect to genomic (nuclear) DNA using primers complementary to three genes encoded in the mitochondrial genome –cytochrome B, D-loop3, and mt-nonNUMT– in conjunction with the nuclear-encoded gene β-actin. Real-time qPCR studies showed that mitochondrial DNA content increased in a time-dependent manner upon Ang II stimulation (Figure 5, panel A).

**Figure 5.**
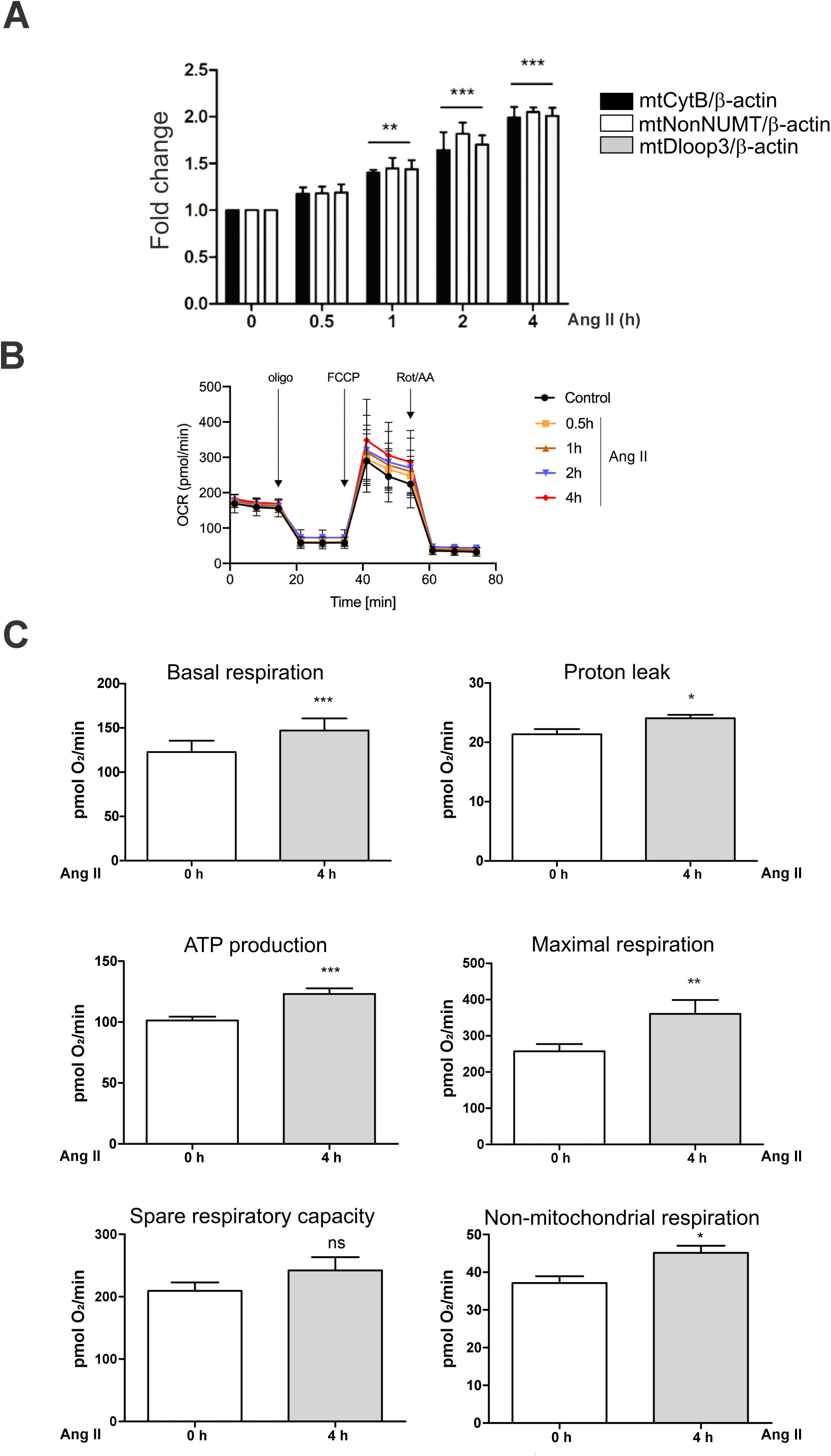
Ang II induces an increase in mitochondrial DNA content, OCR, and respiratory parameters. (A) H295R cells were treated with angiotensin II (Ang II, 100nM) for the times indicated, followed by total DNA isolation. Mitochondrial DNA (mtDNA) content (3 different genes) was quantified by real-time qPCR and normalized to genomic DNA levels (β-actin expression). qPCR data were analyzed using the 2^-ΔΔCt^ method (comparative Ct method) for each experimental sample. Relative genes expression levels (fold change): **p<0.01; *** p<0.001 vs. control cells (Ang II 0 h). For OCR and respiratory parameters measurements, 100,000 cells/well were seeded in 96-well plates in complete medium 48 h prior to the assay. H295R cells were stimulated with angiotensin II (Ang II, 100 nM) for the times indicated. On the day of the assay, the medium was replaced with a specialized Mito Stress Test medium. ATP synthase inhibitor Oligomycin A (2.5 µM) was added to derive ATP-linked OCR, FCCP (0.5 µM) to uncouple the mitochondria for maximal OCR, and Rotenone/Antimycin A (0.5 µM) to inhibit electron transport chain. (B) OCR representative trace in H295R control cells (black circle) and Ang II-stimulated H295R cells (colored, indicated in the graphic). (C) Using OCR data, respiratory parameters were determined and represented in bar graphs: Basal respiration, maximal respiration, spare respiratory capacity, proton leak, ATP-linked respiration, non-mitochondrial respiration. Results are expressed as the mean of pmol O_2_/min ± SD of three independent measurements: ns: non-significant, *p<0.05, **p<0.01 vs control (Ang II, 0 h). Data were analyzed by ANOVA followed by Tukey’s post-hoc test.

### Ang II promotes an increase in mitochondrial function

Given the critical importance of mitochondria in steroid production, we investigated the role of Ang II in bioenergetics, specifically focusing on mitochondrial respiration as a functional parameter.

For this study, control cells and cells stimulated with Ang II for different time periods were analyzed using the XF Cell Mito Stress Test on the Seahorse XFe96 Extracellular Flux Analyzer. OCR quantification allowed the calculation of basal respiration, maximal respiration, non-mitochondrial oxygen consumption, ATP production, proton leak, and spare respiratory capacity in both basal and hormonally stimulated conditions. As assessed in this work, the expression of genes that regulate mitochondrial biogenesis, redox status, and mitochondrial activity is modulated by Ang II. We showed that Ang II promoted an increase in the OCR in a time-dependent manner (Figure 5, panel B).

To further investigate mitochondrial metabolism and function, we calculated the respiratory parameters mentioned earlier based on the OCR measurements. A substantial increase was evident in basal respiration, maximal respiration, ATP production, non-mitochondrial respiration, and proton leakage after 4-hour stimulation with Ang II; nevertheless, no significant differences were found in respiratory reserve capacity (Figure 5, panel C).

### Mitochondrial membrane depolarization regulates Ang II-induced gene expression

In order to determine whether compensatory mechanisms allow H295R adrenocortical cells to counteract mitochondrial depolarization, we incubated H295R cells in the presence of CCCP, a proton ionophore known to induce mitochondrial membrane depolarization. CCCP belongs to a class of lipophilic weak acids that function as proton carriers by selectively increasing the permeability of lipid membranes to protons, thereby dissipating the electrochemical gradient [33].

Our results demonstrate that decrease in mitochondrial activity effectively promotes the upregulation of mRNA levels for various genes which may, in turn, counteract the negative effects of CCCP on mitochondrial function (Figure 6). It is worth noting that 5 µM CCCP, as used in the present study, has previously been reported by our group to significantly inhibit Ang II-stimulated aldosterone production in H295R cells [26]

**Figure 6.**
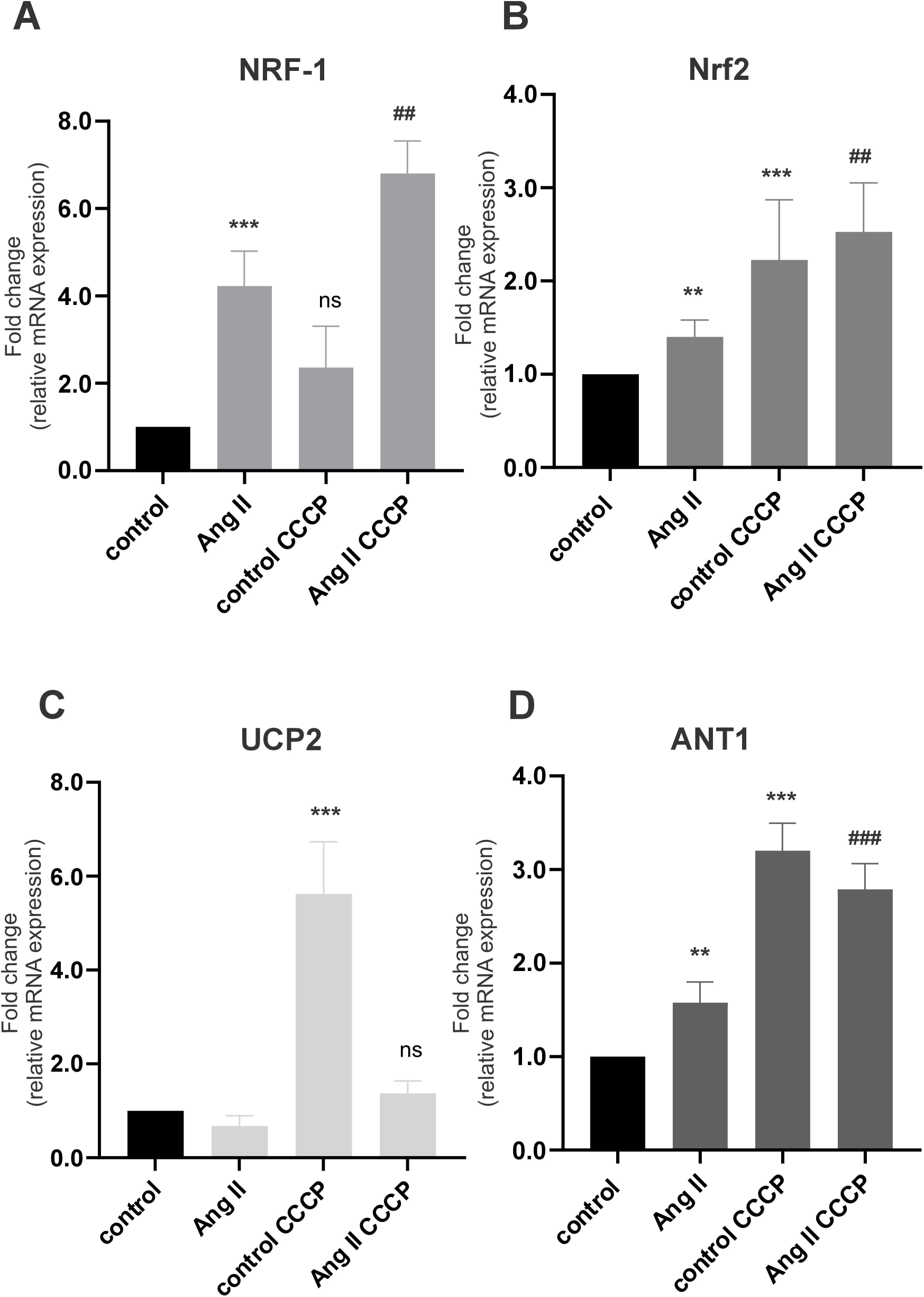
Mitochondrial membrane depolarization modulates the expression of NRF1, Nrf2, UCP2, and ANT1 under both basal and hormone-stimulated conditions. H295R cells were serum-starved for 24 h, pre-incubated for 30 min with CCCP (5 μM) or DMSO (0.1%) as vehicle in the corresponding wells, and subsequently stimulated with angiotensin II (Ang II, 100 nM) for 2 h. Total RNA was isolated and subjected to real-time qPCR using specific primers. Cyclophilin was used as an internal control. qPCR data were analyzed using the 2^-ΔΔCt^ method (comparative Ct method) for each experimental sample. Relative (A) NRF-1, (B) Nrf2, (C) UCP2 and (D) ANT1 expression levels (fold change): ns: non-significant; **p <0.01; ***p<0.001 vs. control (Ang II 0 h); ^##^p <0.01; ^###^p<0.001 without CCCP. Results are expressed as the mean ± SD of three independent experiments. Data were analyzed by ANOVA followed by Tukey’s post-hoc test.

### The cAMP/PKA signaling pathway modulates the expression of genes regulating mitochondrial biogenesis, redox status, and mitochondrial activity

The ACTH signal induces the activation of PKA via an increase in intracellular cAMP, a transduction cascade different from that triggered by Ang II. Given that our cellular model functionally responds to these different stimuli by producing steroids [27], we investigated whether the cAMP/PKA pathway regulates the expression of key mitochondrial biogenesis and activity factors. For this purpose, we used 8-Bromoadenosine 3’,5’-cyclic monophosphate (8Br-cAMP), a cell-permeant cAMP analog that mimics the effect of the endogenous second messenger [28], as H295R cells often exhibit a diminished response to native ACTH due to reduced receptor sensitivity.

The analysis of NRF-1 by Real-time qPCR showed a time-dependent increase in NRF-1 mRNA levels upon 8Br-cAMP treatment, with a marked increase in NRF-1 expression as fast as 30 min after the stimuli (Figure 7, panel A).

**Figure 7.**
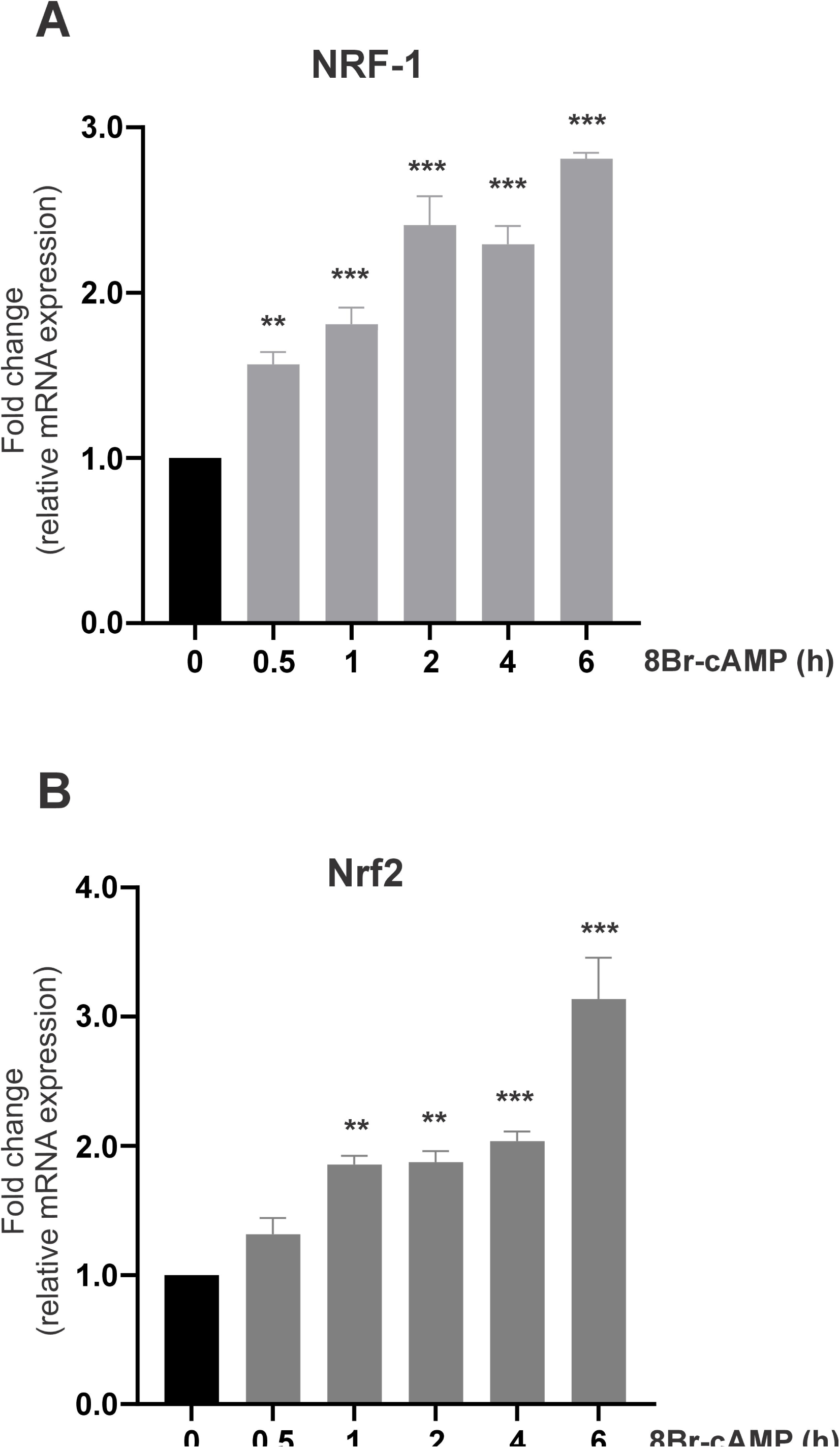
8Br-cAMP modulates NRF-1 and Nrf2 mRNA levels in a time-dependent manner. H295R cells were serum-starved for 24 h and subsequently stimulated with 8Br-cAMP (1 mM) for the times indicated (hours, h). Total RNA was isolated, and real-time qPCR was performed using specific primers. Cyclophilin was used as an internal control. qPCR data were analyzed using the 2^-ΔΔC^ method (comparative Ct method) for each experimental sample. Relative (A) NRF-1 and (B) Nrf2 expression levels (fold change): **p <0.01; *** p<0.001 vs. control cells (Ang II 0 h). Results are expressed as the mean ± SD of three independent experiments. Data were analyzed by ANOVA followed by Tukey’s post-hoc test.

As previously mentioned, Nrf2 has been described as an endogenous antioxidant which prevents excessive ROS production and its deleterious pro-oxidant effects. Incubation of H295R cells with 8Br-cAMP induced an increase in Nrf2 expression up to 4 h of stimulation, followed by an even more pronounced upregulation at 6 h, as shown in Figure 7, panel B.

Next, we examined the regulation of the ETC and OXPHOS activity by the analysis of UCP2 and ANT1. 8Br-cAMP treatment promoted an increase in UCP2 mRNA levels at early time points of stimulation; however, these levels began to decline between 4 and 6 h of exposure (Figure 8, panel A) remaining elevated compared to non-stimulated control cells.

**Figure 8.**
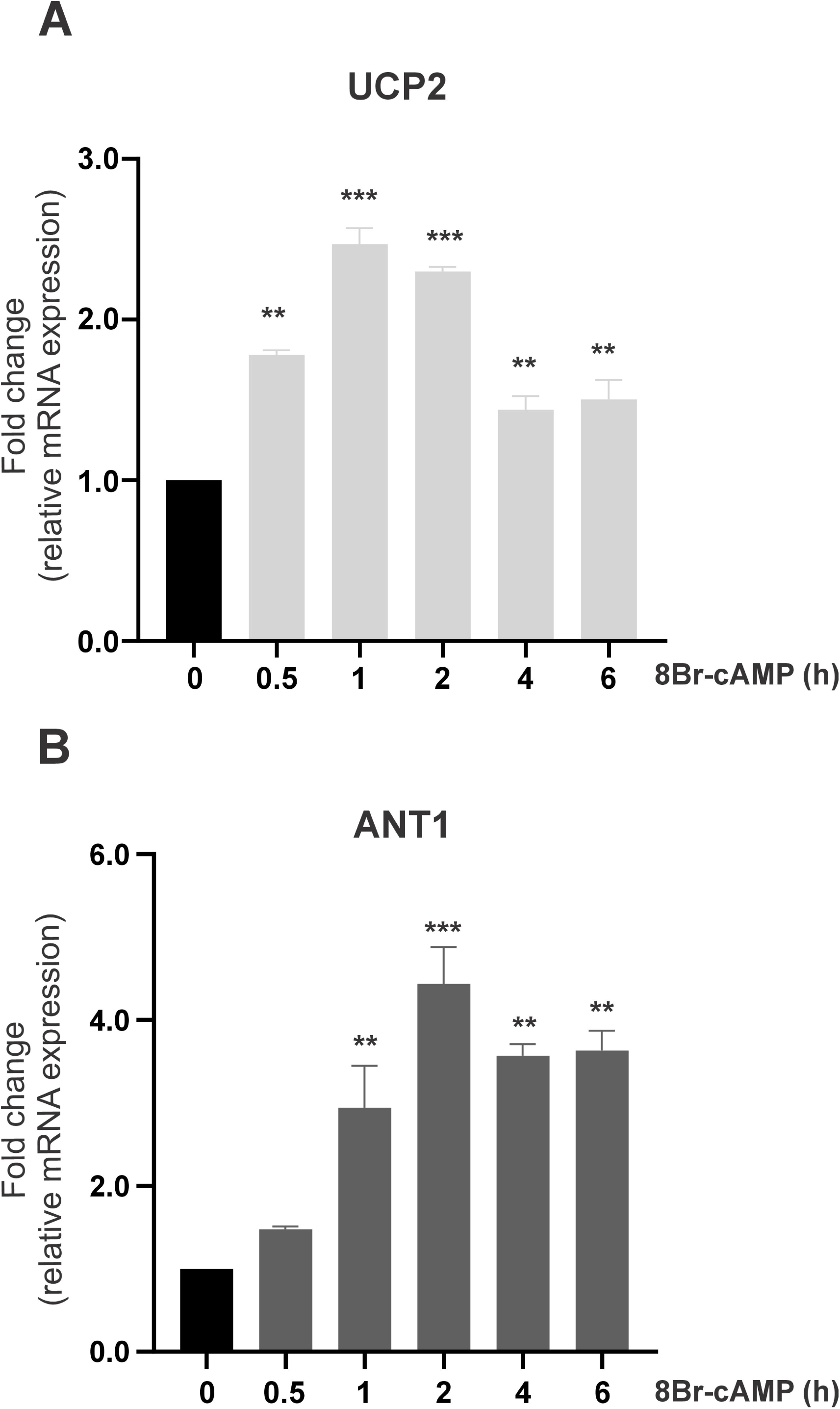
UCP2 and ANT1 expression is regulated by 8Br-cAMP. H295R cells were serum-starved for 24 h and subsequently stimulated with 8Br-cAMP (1mM) for the times indicated (hours, h). Total RNA was isolated and subjected to real-time qPCR using specific primers. Cyclophilin was used as an internal control. qPCR data were analyzed using the 2^−ΔΔCt^ method (comparative Ct method) for each experimental sample. Relative (A) UCP2 and (B) ANT1 expression levels (fold change): **p<0.01; *** p<0.001 vs. control cells (8Br-cAMP 0 h). Results are expressed as the mean ± SD of three independent experiments. Data were analyzed by ANOVA followed by Tukey’s post-hoc test.

We found that stimulation with 8Br-cAMP induced a time-dependent increase in ANT1 expression (Figure 8, panel B), with a peak at 2 h but levels still above those of non-stimulated cells up to 6 h.

To further analyze hormone-triggered pathways in human adrenocortical cells, we evaluated the effect of time-dependent 8Br-cAMP stimulation on mitochondrial activity in H295R cells using MitoTracker fluorescence. We found that the cAMP pathway induced an increase in mitochondrial activity, as evidenced by an increase in red fluorescence intensity up to 2 h of stimulation (Figure 9, panels A–D). This may be driven by the upregulation of proteins such as ANT1, which reaches peak expression at 2 h of 8Br-cAMP treatment. Furthermore, ANT1 can be activated at early time points via ERK1/2 phosphorylation [34]. Indeed, the fact that 8Br-cAMP promotes the rapid phosphorylation and activation of ERK1/2 [31] may be thought of as a plausible mechanism for cAMP-mediated ANT1 activation in H295R cells. We also observed that mitochondrial activity returned to levels near control after 2 h of treatment, which coincides with the downward trend in ANT1 and UCP2 expression. CCCP treatment resulted in a marked decrease in red fluorescence, with quantification showing a reduction in mitochondrial activity following mitochondrial uncoupling (Figure 9, panel G).

**Figure 9.**
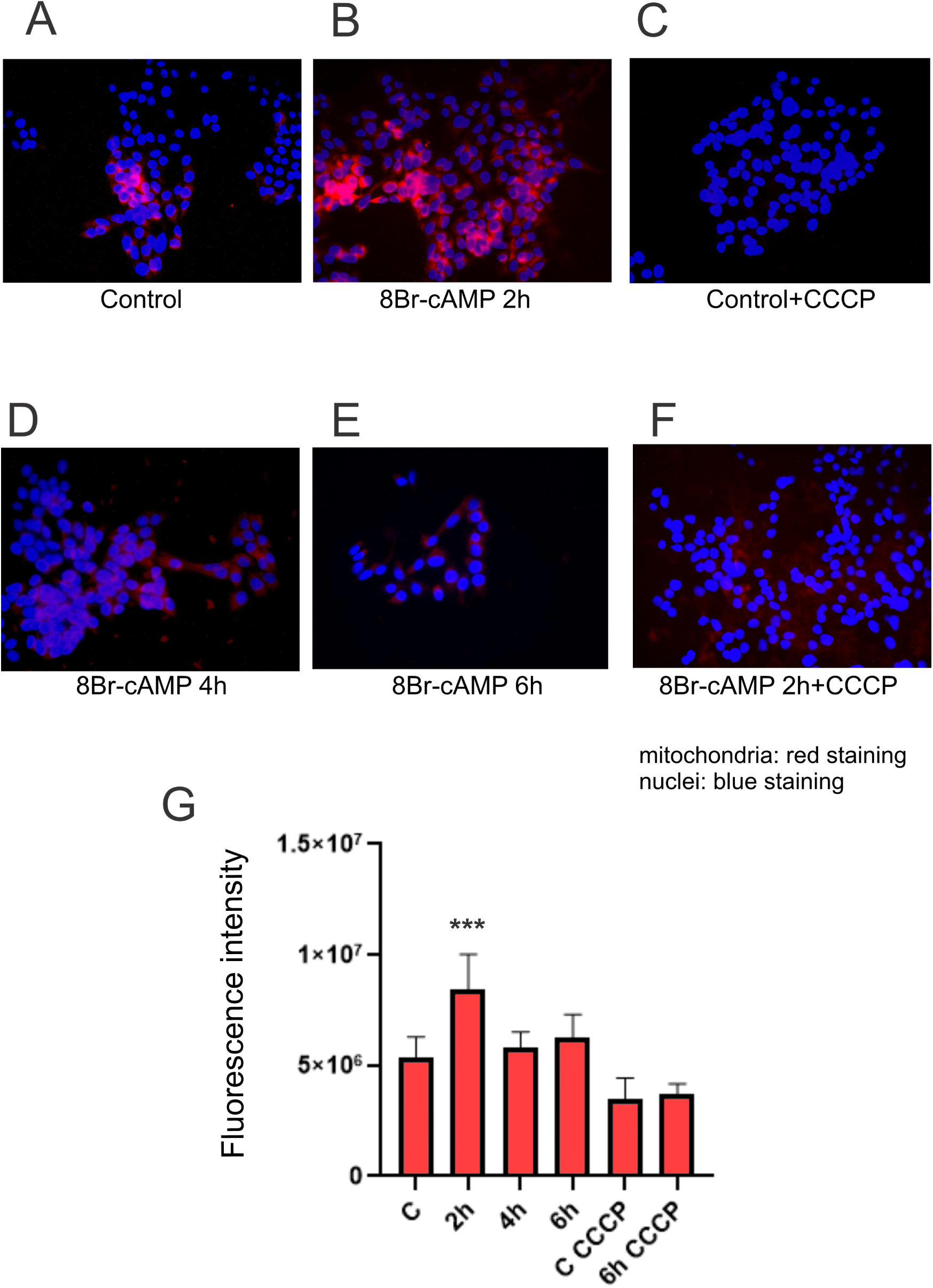
8Br-cAMP stimulation promotes mitochondrial activity. H295R cells were grown on poly-L-lysine glass coverslips, serum-starved for 24 h, and subsequently incubated with CCCP (5 μM) and/or DMSO (0.1%) as vehicle in the corresponding wells (F), followed by stimulation with or without 8Br-cAMP (1 mM) for the times indicated (hours, h) (A-F) (A-D). Afterwards, cells were incubated with 150 nM MitoTracker Red (MTR) at 37°C in fresh medium for 45 min, and samples were processed for analysis using a Nikon fluorescence microscope (40x magnification). (G) In the bar graph, differences in red signal intensity (mitochondrial signal) are expressed as fluorescence intensity (FIJI software) relative to the number of cells within the same ROI (containing 15–20 cells/ROI in all cases). Red (MTR), and ultraviolet (DAPI) channels were studied. Representative images are shown of 10 images analyzed from 3 independent experiments (n=3). Images were obtained in all cases with the same exposure, the background was discarded, and the intensity of the red fluorescent signal was analyzed with FIJI software. ***p<.001 vs control (Ang II 0 h). Scale bar: 20 µm for every image. Data were analyzed by ANOVA followed by Tukey’s post-hoc test.

### Mitochondrial membrane depolarization regulates gene expression induced by cAMP signaling

Next, we assessed the effect of the cAMP/PKA pathway on mitochondrial retrograde signaling. Under basal conditions, the decrease in mitochondrial activity and the concomitant increase in ROS production triggered retrograde signaling and induced the expression of NRF-1, Nrf2, UCP2, and ANT1. This response is likely to promote mitochondrial biogenesis and antioxidant factors while reactivating mitochondrial activity, as shown in Figure 10. After 2 h of stimulation with 8Br-cAMP, we observed an increase in mRNA levels of all four genes, as previously demonstrated. However, pre-incubation with CCCP exerted different effects on gene induction modulated by 8Br-cAMP (Figure 10).

**Figure 10.**
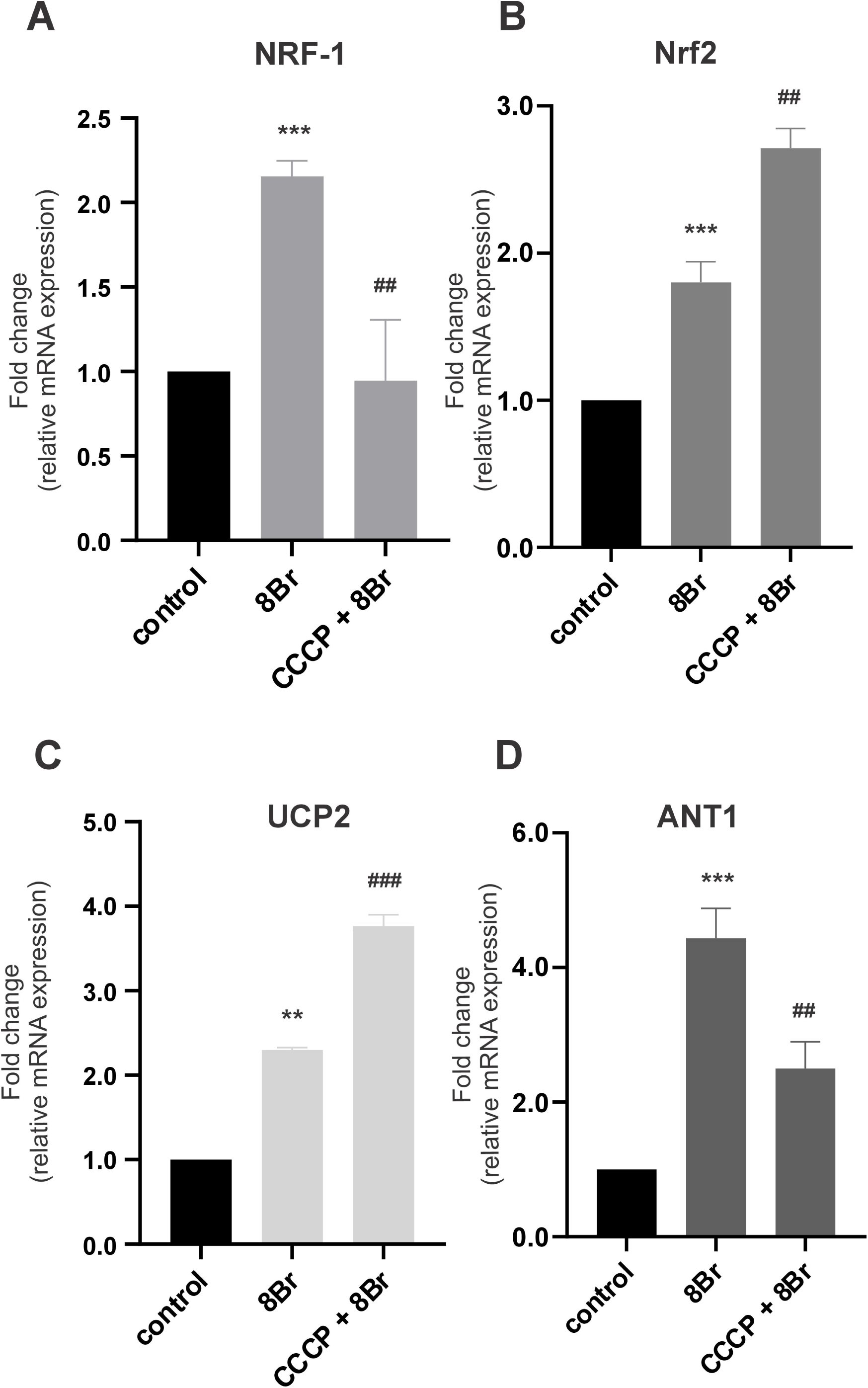
Mitochondrial membrane depolarization modulates the expression of NRF1, Nrf2, UCP2, and ANT1 under 8Br-cAMP stimulation. H295R cells were serum-starved for 24 h, pre-incubated for 30 min with CCCP (5 μM) or DMSO (0.1%) as vehicle in the corresponding wells and subsequently stimulated with 8Br-cAMP (1 mM) for 2 h (indicated as 8Br). Total RNA was isolated and subjected to real-time qPCR using specific primers. Cyclophilin was used as an internal control. qPCR data were analyzed using the 2^-ΔΔCt^ method (comparative Ct method) for each experimental sample. Relative (A) NRF-1, (B) Nrf2, (C) UCP2 and (D) ANT1 expression levels (fold change): **p <0.01; ***p<0.001 vs. control (Ang II 0 h); ^##^p <0.01; ^###^p<0.001 without CCCP. Results are expressed as the mean ± SD of three independent experiments. Data were analyzed by ANOVA followed by Tukey’s post-hoc test.

## Discussion

In the present study, we demonstrate that hormonal stimulation through the Ang II and cAMP signaling pathways coordinately regulates the mitochondrial genetic program and bioenergetic function in human adrenocortical cells. Both stimuli promote the expression of transcription factors involved in mitochondrial biogenesis and antioxidant defense, as well as key proteins that support ETC and OXPHOS normal activity, ultimately enhancing mitochondrial respiration, ATP production, and membrane polarization. Although these pathways share common downstream effects, cAMP elicits a faster mitochondrial response than Ang II, highlighting distinct regulatory kinetics. Furthermore, our findings provide the first evidence that mitochondrial depolarization activates retrograde signaling in human adrenocortical cells, revealing a coordinated mechanism by which hormonal signals optimize mitochondrial function to sustain steroidogenesis. As a summary, Figure 11 depicts a schematic representation of these results.

**Figure 11.**
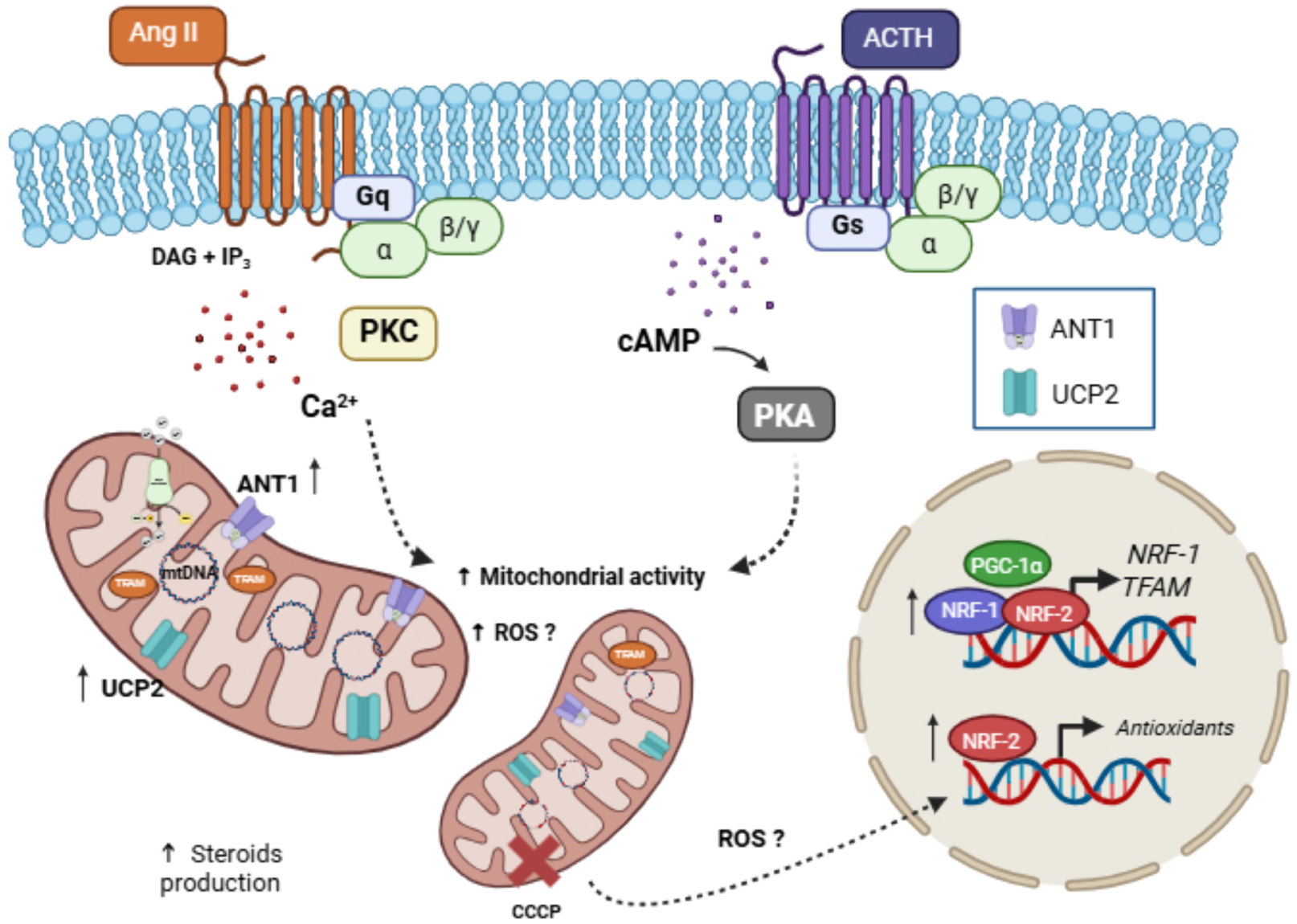
Hormonal regulation of mitochondrial function, biogenesis, and redox homeostasis in H295R adrenocortical cells. Schematic representation of the signaling pathways activated by Ang II and ACTH in H295R human adrenocortical cells and their impact on mitochondrial function. Ang II signaling via Gq promotes Ca²⁺ mobilization and DAG/IP₃ production, leading to PKC activation, whereas ACTH signaling via Gs stimulates cAMP/PKA pathways. These cascades modulate mitochondrial activity, steroidogenesis, and ROS production, while also influencing mitochondrial biogenesis through nuclear transcriptional regulators (NRF-1, TFAM) and mitochondrial activity through mitochondrial factors, as UCP2 and ANT1. These hormonal transduction pathways modulate the mitochondria-nucleus retrograde signaling triggered by mitochondrial membrane potential disruptors. The coordinated response integrates mitochondrial dynamics and antioxidant defenses to maintain cellular homeostasis. Created in BioRender. https://BioRender.com/6cr7t65.

The interaction between Ang II signaling and mitochondrial biogenesis, specifically mediated by NRF-1, exhibits a striking context-dependence with divergent phenotypic effects contingent upon the cellular lineage [35,36]. Although Ang II action may be unfavorable to several tissues, results obtained in H295R human adrenocortical cells evidence a functional profile consistent with a physiological rather than a pathological response. In zona glomerulosa cells, Ang II does not behave as an inducer of cellular stress but as the primary regulator of steroidogenesis, promoting aldosterone synthesis and contributing to the trophic maintenance of the adrenal cortex. Considering that steroid biosynthesis critically depends on mitochondrial function — particularly, adequate respiratory capacity and ATP availability [15]— the observation that Ang II stimulation induces the expression of key mitochondrial biogenesis regulators such as NRF-1 in H295R cells bears clear functional relevance (Figure 11). This induction, in turn, favors the expansion and maintenance of the mitochondrial network required to sustain highly steroidogenic metabolic demand.

Despite the relevance of NRF-1, the specific cascade linking Ang II to this factor in the adrenal cortex has not been fully elucidated yet. It has been established that stimuli such as exercise, hypoxia, and inflammatory processes, in conjunction with receptors including estrogen receptor (ER) and ERRα, modulate NRF-1 expression [37]. In this context, previous findings from our laboratory demonstrate that Ang II increases ERRα levels in a time-dependent manner in H295R cells [38], suggesting that NRF-1 regulation could be mediated by the Ang Type 1 receptor (AT1R)/Ang II signaling pathway. Beyond mitochondrial biogenesis, NRF-1 emerges as a central player in steroidogenesis. In Leydig cells, NRF-1 has been reported to bind to the steroidogenesis acute regulatory protein (StAR) gene promoter, thereby promoting its transcription and increasing testosterone production [39]. The present data suggest that a similar mechanism may underlie aldosterone production in the adrenal gland. Furthermore, the recent description of NRF-1 as a cholesterol sensor via its cholesterol-recognition amino acid consensus (CRAC) domain –which allows it to bind excess cholesterol in the endoplasmic reticulum [40]– has revealed a greater degree of complexity: It is hypothesized that NRF-1 may regulate not only biogenesis but also the availability of the essential lipid substrate for hormone synthesis.

Increased mitochondrial activity inherently entails higher production of ROS. To counteract ROS production, the cellular response includes the activation of Nrf2, the master regulator of the antioxidant response. Nrf2 also modulates mitochondrial function by directly activating NRF-1 and TFAM and controlling the expression of PGC-1α [41]. In relation to steroidogenesis, Nrf2 promotes the increase of StAR and P450scc/CYP11A1 expression and testosterone serum levels in old mice [42] and redox status in mice adrenal gland cells is crucial for CYP11B1 activity [43], inferring a possible role of Nrf2 in this process. In H295R adrenal cells, Ang II signaling has been shown to generate a rapid increase in ROS, which is necessary for CYP11B2 expression and aldosterone production [44]. Our finding that Ang II does not significantly increase Nrf2 levels within the first 30 minutes suggests that the early signaling phase permits a transient rise in ROS required for signal transduction. The subsequent upregulation of Nrf2 may then serve as a protective mechanism to limit oxidative damage associated with sustained ROS production. It has been demonstrated that the PI3K/Akt pathway can orchestrate this process, in agreement with the activation of this cascade in H295R adrenocortical cells by Ang II [25]. This represents a possible mechanism for phosphorylation-mediated Nrf2 activation [45,46].

Uncoupling proteins such as UCP2 and translocases such as ANT1 play a crucial role in dissipating the proton-motive force [47]. UCP2 is expressed in a variety of tissues and has been linked to a protective effect against oxidative damage through a process of mild uncoupling that limits ROS formation [48,49]. Interestingly, it has been described that UCP2 may influence androgen synthesis by affecting CYP11A1 expression in polycystic ovary syndrome patients [50] and UCP2 directly participates in progesterone production in human cumulus cells [51]. Our data demonstrate that Ang II regulates UCP2 in a biphasic manner, allowing an initial window of high ROS production for steroidogenesis, followed by an increase in UCP2 to sustain ETC activity, which in turn helps the continuity of adrenal steroidogenesis. A comparable mechanism has been observed in pancreatic islets, where Ang II, via the AT1R receptor, has been shown to increase ROS levels and UCP2 expression [52]. Concurrently, ANT1 facilitates not only the transportation of nucleotides but also the promotion of proton leak in the presence of fatty acids [53] thereby exerting a cytoprotective effect and regulating mitochondrial permeability [54]. Ang II mediated modulation of UCP2 and ANT1 is shown in Figure 11.

The cAMP pathway has been observed to regulate the functional capacity of respiratory complexes at both the post-translational and transcriptional levels through the action of ACTH [55]. Components such as the PKA-anchoring proteins (AKAPs) and PKA have been associated with mitochondria [56], where CREB phosphorylation has been shown to promote NRF-1 upregulation and OXPHOS activity [57] and thus facilitate mitochondrial biogenesis [58]. In H295R adrenocortical cells, 8Br-cAMP has been shown to induce a significant increase in NRF-2 at 6 hours, possibly to regulate the homeostasis of proteins like StAR, whose degradation by the LON protease –regulated by NRF-2– prevents mitochondrial stress from protein overload [59]. Regarding the effects of cAMP/PKA on oxidative metabolism and redox status, several studies have shown a decrease in ROS levels [60], while others have reported an increase [61]. Nevertheless, the increase in mitochondrial proteins such as UCP2 and ANT1 following cAMP stimulation appears to be an adaptive response to maximize metabolic rate [62] and energy/heat generation [63]. A potential underlying mechanism could involve the induction of ANT1, which facilitates ADP/ATP transport, thereby increasing ETC activity. The temporal regulation of ANT1 by 8Br-cAMP may exert a dual effect on OXPHOS activation; ANT1 increased expression may favor hormone-induced steroidogenesis and further sustained ANT1 levels could be crucial to regulate mitochondrial activity and to avoid excessive ROS production.

Interestingly, the activation profiles of Ang II and cAMP/PKA stimuli differ, with 8Br-cAMP exhibiting faster kinetics than Ang II in the induction of genes regulating mitochondrial function. Although these differences were not explored in depth in the present study, the distinct signaling cascades activated by each stimulus —together with the fact that 8Br-cAMP can permeate the plasma membrane independently of cell surface receptors— may account for the divergent activation profiles observed in the expression of transcription factors and proteins involved in mitochondrial biogenesis and activity.

Disruption of the ΔΨ*m* by the ionophore CCCP [33] triggers a mitochondrial retrograde signaling response [64]. Impairment of mitochondrial function has been demonstrated to lead to Ca^2+^ efflux and the activation of pathways such as CaMK and MAPKs, which induce PGC-1α, NRF-1, and Nrf2 to reactivate biogenesis and antioxidant protection [65].

In this work, we demonstrate that the effect of CCCP increases the expression of NRF-1, Nrf2, and ANT1 in both basal conditions and upon Ang II stimulation. In contrast, UCP2 expression is elevated by CCCP but reaches statistical significance only in basal conditions, with a modest increase observed in Ang II–stimulated cells. Given that UCP2 expression shows a slight decrease after 2 hours of Ang II stimulation, these findings suggest that retrograde signaling triggered by CCCP-induced ΔΨ*m* disruption does not override the effect of Ang II on UCP2 expression. CCCP dissipates the ΔΨ*m*, which —as observed in this work— reduces the 8Br-cAMP–mediated induction of NRF-1 and ANT1, while increasing the expression of Nrf2 and UCP2 stimulated via the cAMP pathway. These results hint at distinct mechanisms in the hormonal regulation of each gene. The cAMP/PKA-mediated induction of NRF-1 described here and reported in other models [57] may be negatively affected by CCCP. Even under depolarization, increased ROS regulate genes like UCP2 and ANT1 to abolish or reduce superoxide production [66]. In addition, CCCP-induced increases in ROS and H₂O₂ levels have been associated with a significant reduction in PGC-1α expression in cultured rat muscle cells [67], a mechanism that may account for the decrease in NRF-1 expression observed in our cellular model.

The increased expression of Nrf2 and UCP2 may function as an antioxidant defense mechanism in response to elevated ROS levels and oxidative stress induced by CCCP (see Figure 11). It is noteworthy that increased ROS levels inhibit Nrf2 proteasomal degradation [68], thereby promoting protein accumulation. Together with the increase observed in mRNA levels, this may reinforce the antioxidant and cytoprotective response to CCCP. Although 5 µM CCCP is widely used to induce ΔΨ*m* depolarization and to investigate retrograde mitochondrial-to-nuclear signaling, it is well established that CCCP can also affect additional cellular pathways. Therefore, we cannot exclude the possibility that some of these pathways contribute to the differential responses to CCCP observed following Ang II and 8Br-cAMP stimulation.

In regard of our bioenergetic analysis our results revealed that Ang II significantly increased OCR, basal and maximal respiration, ATP production, and proton leak. The increase in maximal respiration indicates that H295R cells possess an elevated spare respiratory capacity, which allows them to quickly respond to energy demands.

The present study demonstrates that Ang II stimulation induces significant increases in ATP production, non-mitochondrial respiration –supported by cytoplasmic oxidases–, and proton leak. As OXPHOS is not fully coupled, some protons may leak across the inner mitochondrial membrane independently of mitochondrial ATP synthase.

In relation to the impact of Ang II on mitochondrial metabolism, extensive research has demonstrated that in various tissues, the negative or deleterious effects of Ang II include decreased mitochondrial respiration, ATP synthesis, and maximal respiration [69,70]. Other studies have reported that Ang II promotes the migration of human cardiac fibroblasts by increasing ATP synthesis, maximal respiration, and spare respiratory capacity [71]. To date, no literature has been found linking Ang II stimulation to these mitochondrial parameters in adrenocortical cells. Therefore, taken together, these findings imply that the effect of Ang II on mitochondrial bioenergetics is highly tissue- or context-dependent.

Despite the prevalence of associations with various pathological processes, proton leak has been shown to regulate several physiological processes, particularly in adipose tissue and muscle [72,73]. Consequently, the term “proton leak” is used to denote the residual basal respiration that remains uncoupled from ATP production. Basal proton leak has been described as corresponding to the activity of ANT1 in the presence of fatty acids [74], while stimulated proton leak is primarily due to the expression of UCPs [75,76]. The fact that Ang II promotes the upregulation of both UCP2 and ANT1 suggests a potential mechanism for the increase observed in proton leak. Therefore, in adrenal gland cells—where mitochondrial activity is key in the acute phase of steroidogenesis— Ang II promotes a comprehensive increase in respiratory parameters and mitochondrial functionality. As depicted in Figure 11, hormonal stimulation may be thought to orchestrate mitochondrial biogenesis and enhance metabolic activity, serving as a critical mechanism to initiate and sustain both steroidogenesis and cellular homeostasis within the adrenal gland.

In conclusion, we demonstrate for the first time that distinct hormonal signals modulate mitochondrial functionality in H295R human adrenocortical cells, which may positively impact the essential steroidogenic function of this cell type. These findings are in agreement with the strong association between mitochondrial dysfunction and the pathophysiology of several adrenal diseases.

## Data Availability

Some or all datasets generated during and/or analyzed during the current study are not publicly available but are available from the corresponding author on reasonable request.

## Funding

This work was supported by the National Scientific and Technical Research Council (CONICET) – Argentina, P.I.P.: (2021-2023, 11220200101892CO to C. Poderoso), University of Buenos Aires – Argentina (UBACYT 2020: 20020190100040BA, to P.M. Maloberti and C. Poderoso), National Agency for Scientific and Technological Promotion, (former MinCyT), (PICT-2018-02254 to C. Poderoso). The funders had no role in studying design, data collection and analysis, decision to publish, or preparation of the manuscript.

## Abbreviations

Ang II: angiotensin II
ANOVA: analysis of variance
ANT: adenine nucleotide translocase
ATP: adenosine-5′-triphosphate
AT1R: angiotensin II type 1 receptor
BSA: bovine serum albumin
cAMP: cyclic-adenosine monophosphate
ER: endoplasmic reticulum
ETC: electron transport chain
FCCP: carbonyl cyanide p-trifluoromethoxyphenylhydrazone
mtDNA: mitochondrial DNA
NRF-1: Nuclear respiratory factor type 1
Nrf2: nuclear factor erythroid 2-related factor 2
OCR: oxygen consumption rate
OXPHOS: oxidative phosphorylation
PBS: phosphate-buffered saline
PKA: protein kinase A
PKC: protein kinase C
SDHA: succinate dehydrogenase subunit A
ROI: region of interest
UCP2: uncoupling protein 2
ΔΨ*m*: mitochondrial membrane potential

## Acknowledgments

We are very thankful to Prof. Dr. Thomas Langer’s Laboratory members at Max Planck Institute (MPI) for Biology of Aging, Cologne, Germany, for assisting with the SeaHorse experiments and further analysis.

We thank María M. Rancez for providing language help and writing assistance with the manuscript.

## Notes

### Competing Interest Statement

The authors have declared no competing interest.

### Summary of Updates

The manuscript has been extensively revised to better focus on the main ideas, and several figures have been revised for greater clarity.

